# Chromosome-wide distribution and characterization of intersubspecific meiotic noncrossovers in mice

**DOI:** 10.1101/792226

**Authors:** Vaclav Gergelits, Emil Parvanov, Petr Simecek, Jiri Forejt

## Abstract

During meiosis, the recombination-initiating DNA double-strand breaks (DSBs) are repaired by crossovers or noncrossovers (gene conversions). While crossovers are easily detectable, the noncrossover identification is hampered by the small size of their converted tracts and the necessity of sequence polymorphism to detect them. We report identification and characterization of a mouse chromosome-wide set of noncrossovers by NGS sequencing of 10 mouse chromosome substitution strains. Based on 94 identified noncrossovers we determined the mean length of a conversion tract to be 32 basepairs. The spatial chromosome-wide distribution of noncrossovers and crossovers significantly differed, though both sets overlapped the known hotspots of PRDM9-directed histone methylation and DNA DSBs, thus proving their origin in the standard DSB repair pathway. A significant deficit of noncrossovers descending from asymmetric DSBs pointed to sister chromatids as an alternative template for their repair. The finding has implications for the molecular mechanism of hybrid sterility in mice.

## Introduction

Homologous recombination between paternal and maternal chromosomes and their synapsis are unique features of the meiotic phase of gametogenesis in the majority of the sexually reproducing organisms. In humans, mice and most other mammals the localization of recombination breakpoints is under the control of the PRDM9 histone methyltransferase. The PRDM9 zinc finger domain binds at allele-specific sites of genomic DNA and its PR/SET domain ensures trimethylation of histone 3 at lysine 4 and lysine 36 (H3K4me3 and H3K36me3) surrounding both sides of the binding motif^1–4^. The SPO11 topoisomerase-like protein targets a subset of trimethylated PRDM9 binding sites and generates programmed DNA double-strand breaks (DSBs)^5,6^. The DSBs are repaired by homologous recombination, either by crossovers (COs) accompanied by the exchange of chromatid arms between homologous chromosomes or by noncrossovers (NCOs, gene conversions) created by unidirectional transfer of genetic information from the homologous chromatids. Conversion tracts, usually longer than in NCOs, also accompany COs^7^. The COs are under a strong homeostatic control ensuring at least one CO per chromosome arm (obligate crossover rule)^8,9^, a condition necessary for the proper segregation of homologous chromosomes into gametes. In contrast, NCOs are indispensable for synapsis of homologous chromosomes as illustrated by asynapsis in spermatocytes with reduced or absent DSBs due to attenuated or abolished expression of the *Spo11* gene^10–13^. The significance of NCOs for meiotic synapsis of homologous chromosomes is further corroborated by the synapsis failure in meiocytes of the mouse subspecific hybrids with a reduced number of repairable (symmetric) DSBs^14–16^. The NCO events are hard to detect because the conversion tracts are rather short, estimated at 86 ± 49 bp in two mouse hotspots^17^ and 55-290 bp in humans^18^. Moreover, the NCOs are detectable only when the conversion tracts carry SNPs or short indels distinguishing the paternal from the maternal sequence. Thus, the efficiency of identification of NCOs depends on the density and distribution of these allelic variants.

To identify the NCOs and their chromosome-wide distribution we took advantage of the fact that NCOs are enriched in consomic chromosomes of chromosome substitution (consomic) strains. To construct a consomic strain only the individuals with nonrecombinant donor chromosomes (without COs) are selected for transfer to the next generation during 10 consecutive backcrosses to the recipient inbred strain^19,20^ (Figure 1). Hence the transferred chromosome is 10-fold enriched for NCOs but devoid of COs. In the final intercross the consomic chromosome is fixed in the homozygous state allowing direct identification of NCOs by sequencing. Here we searched for NCOs in 10 strains of a panel of C57BL/6J-Chr #^PWD^/Ph/ForeJ (abbreviated B6.PWD-Chr #) mouse consomic strains^20^. In these mice the donor chromosome descends from the PWD/Ph (PWD) inbred strain derived from the *Mus musculus musculus (M. m. musculus)* subspecies^21^, while the genome of the recipient C57BL/6J (B6) strain is predominantly of *Mus musculus domesticus (M. m. domesticus)* origin^22^. Because both subspecies diverged roughly 350-500 thousand years ago^23,24^ the high level of their genome disparity, one SNP per 130 bp, is comparable to the divergence between the human and chimpanzee^25^. The accumulation of NCOs during the backcrosses and the higher chance to identify them by converted SNPs are the factors that considerably amplified the efficiency of NCO mapping. Moreover, better characterization of NCOs can contribute to genetic dissection of reproductive isolation between the mouse subspecies. The difficulties to repair DNA DSBs by homologous recombination and subsequent deficiency of NCOs was proposed to interfere with the synapsis of homologous chromosomes and cause early meiotic arrest in the (PWD × B6)F1 model of hybrid male sterility^14–16,26,27^.

**Figure 1.**
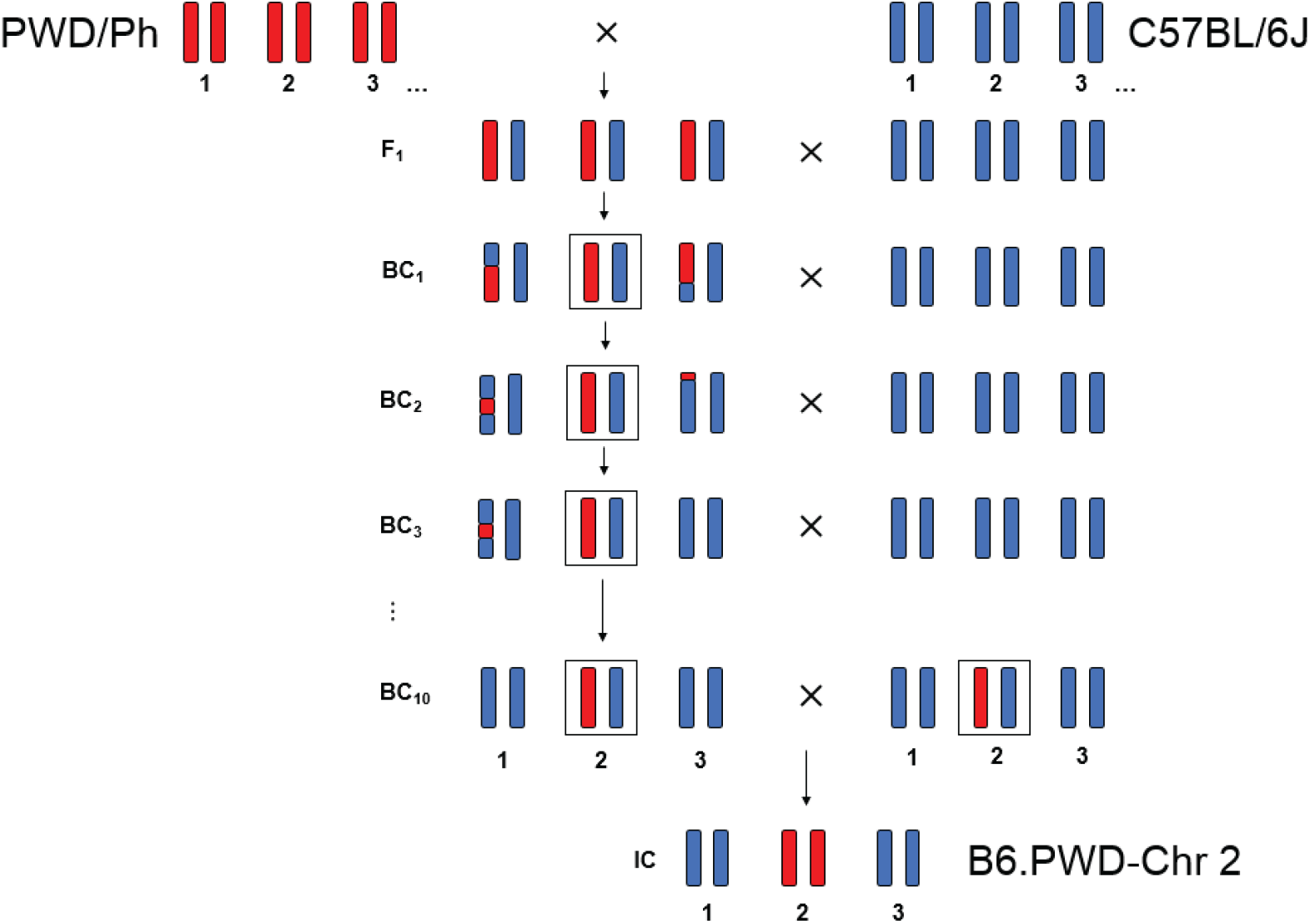
Scheme of construction of chromosomal substitution strains B6.PWD-Chr #.The donor chromosomes (PWD) were transferred to the recipient inbred strain (B6) by ten consecutive backcrosses. In the final intercross the consomic chromosome was fixed in homozygous state allowing direct identification of NCOs by sequencing. The example of B6.PWD-Chr 2 is shown here.

## Results

### Chromosome-wide identification of noncrossovers

To identify NCOs we sought the B6-PWD SNPs in the PWD sequence of individual consomic chromosomes (Figure 2a). Nine B6.PWD-Chr # consomic strains carrying PWD autosomes # 1, 2, 3, 4, 5, 6, 17, 18, 19 and B6.PWD-Chr X.1s subconsomic strain with the proximal part of Chr X of PWD origin^28^ were sequenced with the minimal coverage of 24×. The B6 SNPs were detected by simultaneous alignment of the whole genome sequence (WGS) Illumina reads of a consomic sequence to the B6 reference genome, the pseudo-reference of PWK/PhJ^29^, and were compared to WGS of the PWD/PhJ and PWK/PhJ strains. The latter two are closely related *M. m. musculus*-derived inbred strains. The false-positive variants were eliminated by a multi-step filtering procedure based on the comparison of PWD-B6 SNPs in consomic strains and their B6 and PWD progenitors (see Materials and Methods). Altogether we detected 91 NCOs in nine consomic autosomes and three NCOs in the proximal 69.6 Mb of Chr X (Figure 2b, Table 1). Sixty-nine percent of NCO gene conversion tracts encompassed one converted SNP; 20% and 11% carried two and more SNPs (or possibly short indels), respectively.

**Table 1.** Localization and origin of NCO events.

| NCO localization |  |  |  |  | DMC1 coloc.<sup>1</sup> |  |  |  |  | NCO localization |  |  |  |  | DMC1 coloc.<sup>1</sup> |  |  |  |  |
| --- | --- | --- | --- | --- | --- | --- | --- | --- | --- | --- | --- | --- | --- | --- | --- | --- | --- | --- | --- |
| Chr | Start | End | Min<sup>2</sup> | Max<sup>2</sup> |  |  |  |  |  | Chr | Start | End | Min<sup>2</sup> | Max<sup>2</sup> |  |  |  |  |  |
|  |  |  |  |  | B6 | (PWD×B6)F1 | PWD | B6.Prdm9<sup>-/-</sup> | Nonparental<sup>3</sup> |  |  |  |  |  | B6 | (PWD×B6)F1 | PWD | B6.Prdm9<sup>-/-</sup> | Nonparental<sup>3</sup> |
| 1 | 13845860 | 13845929 | 70 | 277 | T | T | F | F | F | 4 | 92481179 | 92481179 | 1 | 596 | F | F | F | F | NA |
| 1 | 46907980 | 46907980 | 1 | 419 | T | T | F | F | F | 4 | 125451332 | 125451332 | 1 | 661 | T | T | F | F | F |
| 1 | 59359320 | 59359321 | 2 | 692 | T | T | F | F | F | 4 | 142990293 | 142990308 | 16 | 364 | T | T | F | F | F |
| 1 | 61872329 | 61872329 | 1 | 69 | T | T | F | F | F | 5 | 6242093 | 6242323 | 231 | 407 | T | T | F | F | F |
| 1 | 63879266 | 63879271 | 6 | 117 | F | T | F | F | T | 5 | 20789067 | 20789067 | 1 | 115 | T | T | F | F | F |
| 1 | 67960742 | 67963400 | 2659 | 3994 | T | T | F | F | F | 5 | 27852657 | 27852657 | 1 | 338 | T | T | F | F | F |
| 1 | 128517316 | 128517318 | 3 | 985 | T | T | F | F | F | 5 | 51691842 | 51691842 | 1 | 92 | T | T | F | F | F |
| 1 | 137135844 | 137135844 | 1 | 141 | T | T | F | F | F | 5 | 55484917 | 55484946 | 30 | 278 | T | T | F | F | F |
| 1 | 165896307 | 165896311 | 5 | 680 | T | T | F | F | F | 5 | 90775650 | 90775650 | 1 | 61 | T | T | F | F | F |
| 1 | 183578162 | 183578162 | 1 | 542 | T | T | F | F | F | 5 | 96571457 | 96571457 | 1 | 94 | T | T | F | F | F |
| 2 | 3736957 | 3736957 | 1 | 57 | T | T | F | F | F | 5 | 102062273 | 102062273 | 1 | 190 | T | T | F | F | F |
| 2 | 5040747 | 5040747 | 1 | 133 | T | T | F | F | F | 5 | 104084656 | 104084656 | 1 | 340 | T | T | F | F | F |
| 2 | 10336587 | 10336592 | 6 | 264 | T | T | F | F | F | 5 | 108572496 | 108572496 | 1 | 441 | T | T | F | F | F |
| 2 | 11688991 | 11689010 | 20 | 549 | F | T | F | F | T | 5 | 117528325 | 117528325 | 1 | 380 | T | T | F | F | F |
| 2 | 13697711 | 13697711 | 1 | 431 | F | T | T | F | F | 5 | 119304743 | 119304775 | 33 | 110 | T | T | F | F | F |
| 2 | 28188305 | 28188305 | 1 | 361 | F | T | F | F | T | 5 | 125695123 | 125695137 | 15 | 211 | T | T | F | F | F |
| 2 | 32226478 | 32226493 | 16 | 147 | F | T | T | F | F | 5 | 141144023 | 141144023 | 1 | 246 | T | T | T | F | F |
| 2 | 58517611 | 58517618 | 8 | 270 | T | T | F | F | F | 5 | 148185214 | 148185214 | 1 | 54 | F | F | F | F | NA |
| 2 | 61489637 | 61489637 | 1 | 643 | T | T | F | F | F | 6 | 18095110 | 18095110 | 1 | 159 | T | T | F | F | F |
| 2 | 72241857 | 72241857 | 1 | 462 | T | T | F | F | F | 6 | 36814629 | 36814629 | 1 | 299 | T | T | F | F | F |
| 2 | 74937052 | 74937064 | 13 | 384 | T | T | F | F | F | 6 | 39355424 | 39355448 | 25 | 176 | F | T | T | F | F |
| 2 | 78357403 | 78357403 | 1 | 50 | T | T | F | F | F | 6 | 101509677 | 101509677 | 1 | 148 | T | T | F | F | F |
| 2 | 79679676 | 79679676 | 1 | 330 | T | T | F | F | F | 6 | 103949614 | 103949614 | 1 | 122 | T | T | T | F | F |
| 2 | 85528071 | 85528071 | 1 | 131 | T | T | F | F | F | 6 | 115864297 | 115864297 | 1 | 315 | F | T | T | F | F |
| 2 | 96760589 | 96760589 | 1 | 169 | T | T | F | F | F | 6 | 128995778 | 128995882 | 105 | 337 | T | T | F | F | F |
| 2 | 116516318 | 116516318 | 1 | 126 | F | T | F | F | T | 6 | 134249703 | 134249745 | 43 | 388 | T | T | F | T | D<sup>4</sup> |
| 2 | 135984537 | 135984537 | 1 | 46 | F | T | F | F | T | 17 | 14350774 | 14350820 | 47 | 587 | T | T | F | F | F |
| 2 | 147805390 | 147805390 | 1 | 383 | T | T | F | F | F | 17 | 25991437 | 25991437 | 1 | 162 | F | T | F | F | T |
| 2 | 150590002 | 150590002 | 1 | 687 | T | T | F | F | F | 17 | 31984106 | 31984106 | 1 | 113 | F | T | T | F | F |
| 2 | 166617293 | 166617524 | 232 | 370 | F | T | F | F | T | 17 | 51576043 | 51576050 | 8 | 491 | F | T | F | F | T |
| 3 | 16828944 | 16828944 | 1 | 267 | T | T | F | F | F | 17 | 67796579 | 67796615 | 37 | 97 | F | F | F | F | NA |
| 3 | 22128566 | 22128566 | 1 | 378 | T | F | F | F | NA | 17 | 79667734 | 79667734 | 1 | 258 | T | T | T | T | D<sup>4</sup> |
| 3 | 36830205 | 36830205 | 1 | 492 | T | T | F | F | F | 18 | 4596959 | 4596972 | 14 | 394 | T | T | F | F | F |
| 3 | 51188409 | 51188409 | 1 | 182 | T | T | F | F | F | 18 | 28878184 | 28878184 | 1 | 305 | T | F | F | F | NA |
| 3 | 65603278 | 65603278 | 1 | 662 | T | T | F | F | F | 18 | 32083435 | 32083435 | 1 | 46 | T | T | F | F | F |
| 3 | 65655136 | 65655136 | 1 | 466 | F | T | F | F | T | 18 | 38278002 | 38278002 | 1 | 251 | T | T | F | F | F |
| 3 | 69739785 | 69739785 | 1 | 232 | T | T | F | F | F | 18 | 40885813 | 40885839 | 27 | 230 | T | T | F | F | F |
| 3 | 85792585 | 85792585 | 1 | 138 | T | T | F | F | F | 18 | 41946723 | 41946739 | 17 | 162 | T | T | F | F | F |
| 3 | 95869211 | 95869211 | 1 | 36 | T | T | F | F | F | 18 | 78715451 | 78715451 | 1 | 72 | F | T | F | F | T |
| 3 | 107617094 | 107617095 | 2 | 185 | T | T | F | F | F | 19 | 4199635 | 4199635 | 1 | 183 | T | T | F | F | F |
| 3 | 130702831 | 130702831 | 1 | 177 | T | T | F | F | F | 19 | 16998839 | 16998847 | 9 | 130 | F | T | F | F | T |
| 3 | 136064382 | 136064385 | 4 | 622 | F | T | F | F | T | 19 | 28429944 | 28429944 | 1 | 112 | T | T | F | F | F |
| 4 | 6396524 | 6396541 | 18 | 399 | F | T | F | F | T | 19 | 28519574 | 28519585 | 12 | 97 | T | T | F | F | F |
| 4 | 10936243 | 10936243 | 1 | 448 | F | T | F | F | T | 19 | 44938647 | 44938648 | 2 | 444 | T | T | F | F | F |
| 4 | 49528333 | 49528367 | 35 | 225 | T | T | F | F | F | X | 42581841 | 42581841 | 1 | 239 | T | T | F | F | F |
| 4 | 59369670 | 59369670 | 1 | 286 | T | T | F | F | F | X | 52427650 | 52427650 | 1 | 220 | F | T | F | F | T |
| 4 | 83240097 | 83240097 | 1 | 260 | F | T | F | F | T | X | 63297750 | 63297750 | 1 | 193 | F | F | F | F | NA |

<sup>1</sup> DMC1 colocalization; data from Davies *et al.* 2016 and Smagulova *et al.* 2016, overlap within ±1000 bp from DMC1 center. T, True; F, False.
<sup>2</sup> Minimal and Maximal NCO conversion tract lengths.
<sup>3</sup> The nonparental event is defined as found in (B6×PWD)F1 but not in parental B6 nor PWD DMC1 datasets.
<sup>4</sup> Default hotspot; i.e. present in B6.Prdm9<sup>-/-</sup> DMC1 dataset.
\* Complex NCO event, see the main text.

**Figure 2.**
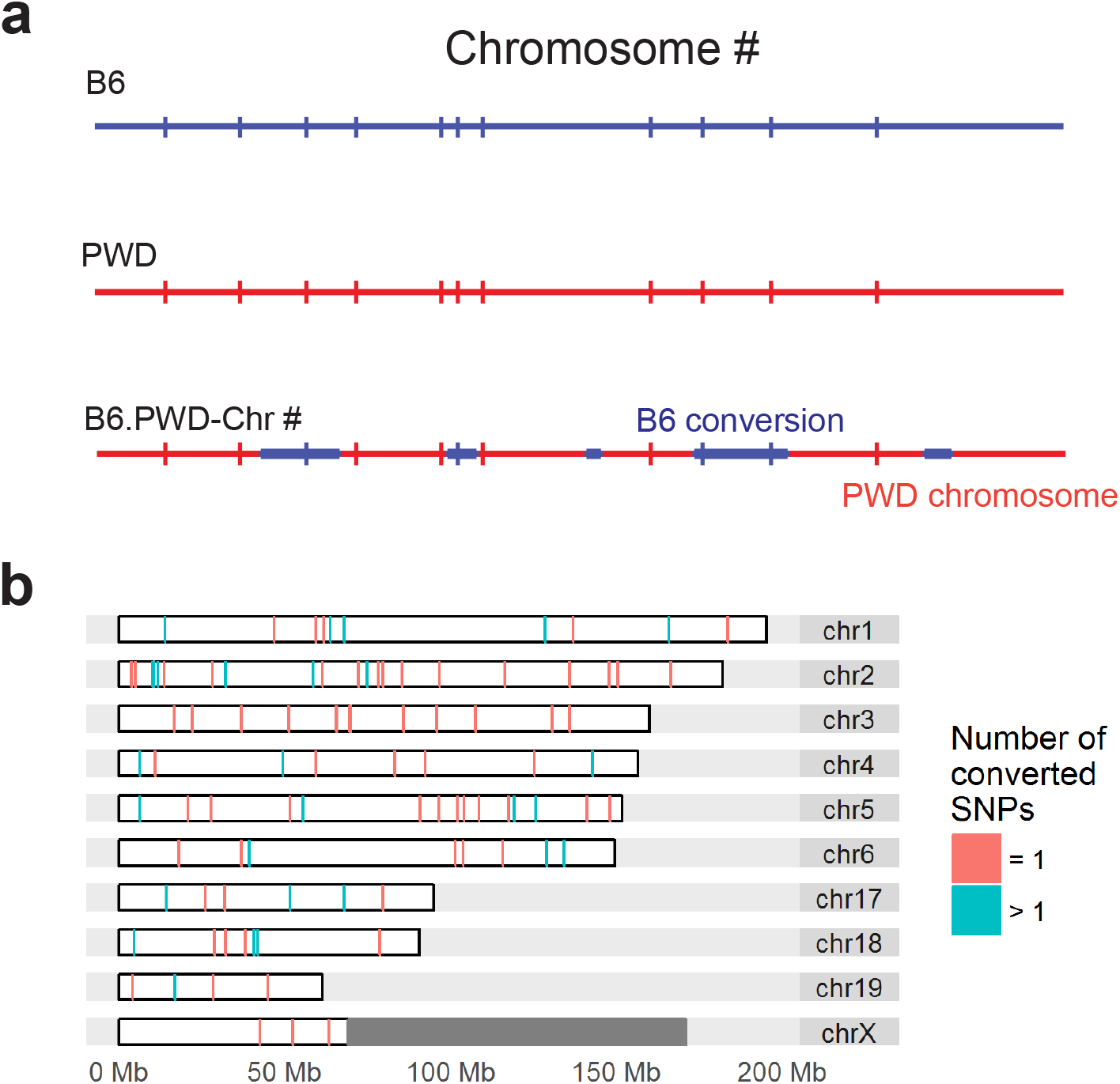
Detection of NCO events. **a** The scheme of NCO detection by comparison of chromosomal substitution strains B6.PWD-Chr# and parental B6 and PWD strains. The NCO site could be covered by 0, 1, or 2 or more SNPs. NCO sites overlapped by 0 SNPs cannot be detected. **b** Localization of NCO events identified on the transferred PWD chromosomes in ten chromosomal substitution strains. The NCO gene conversion with single and multiple converted SNPs are distinguished by color. For the detection of NCOs on chromosome X the subconsomic B6.PWD-ChrX.2 strain carrying the PWD sequence in the proximal 69.6 Mbs was used.

The ranges of possible length of each NCO conversion tract (Table 1) were based on the converted and surrounding non-converted SNPs. The conversion tracts varied in length. For instance, we found 13 tracts necessarily shorter and four tracts necessarily longer than 100 bp (Figure 3). Assuming exponential distribution, we estimated the mean length of the detected NCO conversion tracts at 32 bp. In addition, a single complex conversion event (1%) on Chr 2 contained two converted SNPs 231 bp apart from each other with two non-converted SNPs in between. We confirmed by Sanger sequencing that this complex event more likely resulted from a single meiosis rather than from aggregation of two events in different backcross generations. Thus, the incidence of complex NCO gene conversions was much lower than 46% or 65% reported in the human study^30^.

**Figure 3.**
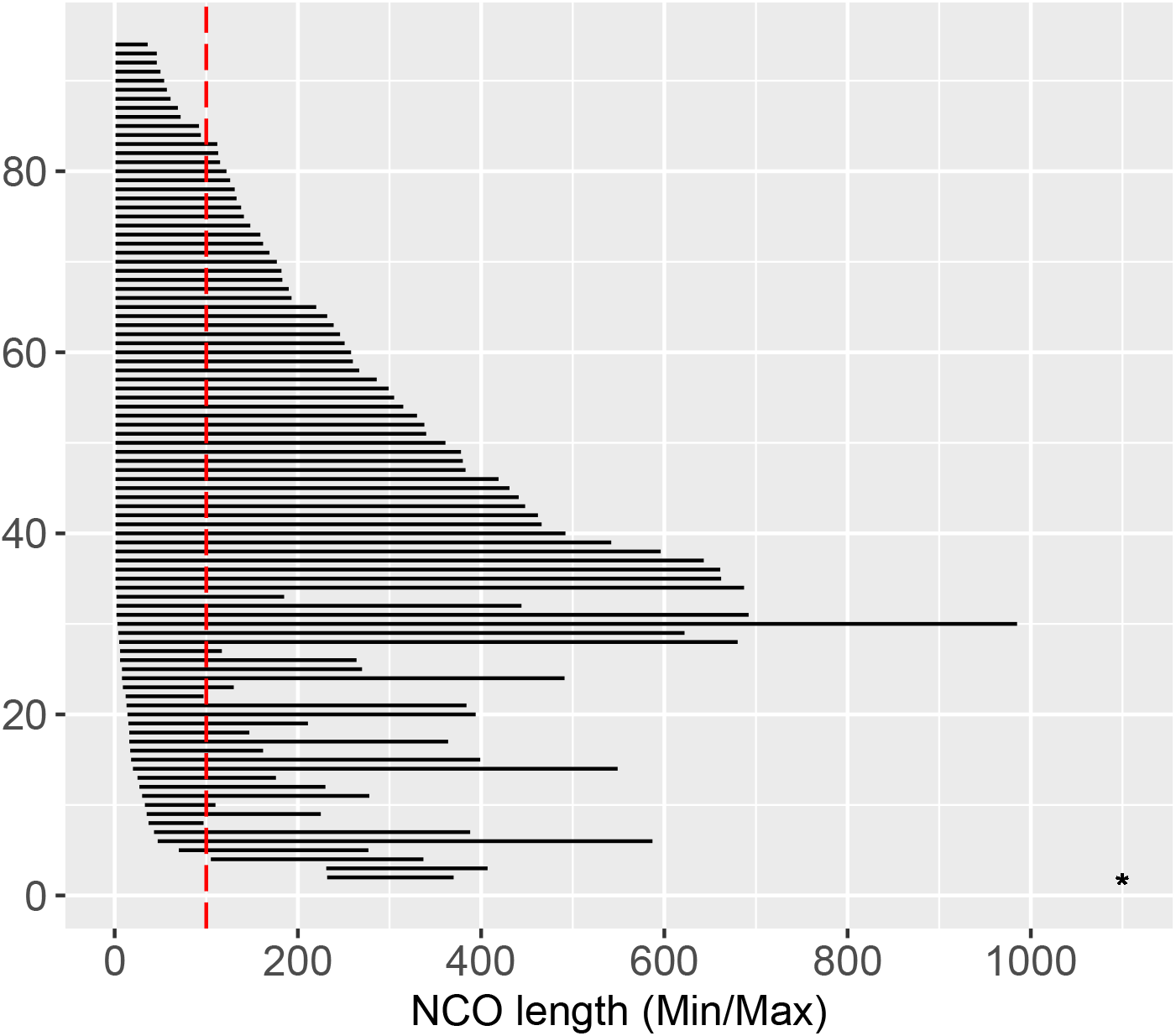
Maximal and minimal possible length of each NCO gene conversion tract. The maximal length is given by the distance of the two nearest non-converted PWD-B6 SNPs. The minimal length is given by the distance of the most outer converted PWD-B6 SNPs; in 71% cases, the conversion tract contains only one SNP giving a minimal length of 1 bp. *The gene conversion with minimal length = 2659 bp and maximal length = 3966 bp is not displayed.

A representative subset (48/94) of identified autosomal NCOs was further validated by Sanger sequencing and all of them were confirmed to be true, proving the 100% specificity of the filtering methods.

### NCOs colocalize with the sites of meiotic DSBs

The meiotic recombination pathway is initiated by activation of PRDM9 binding sites, detectable by chromatin immunoprecipitation of histone 3 trimethylated at lysine 4 and lysine 36 (H3K4me3 and H3K36me3 ChIP-seq) and massively parallel DNA sequencing. About 4700 such sites are activated per leptotene spermatocyte^31^, but only 250-300 acquire DNA DSBs after being targeted by the SPO11 protein. Hotspots of DNA DSBs can be localized and quantified by sequencing SPO11-covalently bound oligos^32^ or by ChIP-seq of DMC1 protein bound DNA recombination intermediates^33^. At kilobase resolution, 96% of the identified NCOs colocalized with the sites of meiotic DMC1 hotspots based on the available data^15,34^ on males of one or more of the strains/crosses – B6, PWD, (B6 × PWD)F1 and (PWD × B6)F1 (Table 1). The overlap showed high specificity of our detection approach, since the random overlap of NCOs with the superset of these DSBs sites was only 7%, thus confirming that the detected NCOs were real. As expected, the same 96% of NCO conversion tracts colocalized with the PRDM9-induced H3K4me3 histone modifications (Table 1-table supplement 1). The NCO counts per chromosome correlated well (Spearman’s ρ = 0.90377, P = 0.0008274) with the SPO11 oligo signal^35^ used as a proxy for the DSB counts (Table 1-table supplement 2), reflecting their dependence as shown in detail below. Information for the H3K36me3 and SPO11 oligo hotspots is not available for (PWD × B6) and (B6 × PWD)F1 hybrids; nevertheless in the subset of NCO sites overlapping the DMC1 hotspots of the B6 strain (72%, 68/94) we assessed the overlap with the PRDM9-mediated H3K36me3 hotspots and the overlap with SPO11 sites of programmed DSBs in B6 meiocytes^35^. NCOs overlapped with H3K36me3 in 79% of cases (all DMC1 sites overlapped H3K36me3 sites in 56%) and NCOs overlapped with SPO11 in 96% cases (all DMC1 sites overlapped SPO11 sites in 72% cases). The higher overlap of NCOs with H3K36me3 and SPO11 sites could be explained by the fact that NCOs are typically localized in the hotter DMC1 sites more likely reflecting higher affinity for SPO11 and H3K36me3. The high overlap of NCOs with the markers of meiotic DSB induction and repair showed that the vast majority of NCOs resulted from the standard meiotic DSB repair pathway. The four DMC1-negative NCO sites, non-overlapping any DSB site in DMC1 datasets of (B6 × PWD)F1, (PWD × B6)F1, B6 and PWD were confirmed by Sanger sequencing to be converted.

### Identification of PRDM9 binding motifs within NCO sites

We identified the PRDM9 binding motif within an interval 500 bp upstream or downstream of the converted SNP in most of the NCOs conversion tracts. When the NCOs were compared with the DMC1 hotspot data, 78% matched with the PRDM9^B6^ hotspots, 5% with PRDM9^PWD^, 11% possibly originated from either PRDM9^B6^ or PRDM9^PWD^ but could not be determined, 2% were default (present in the B6.*Prdm9*^−/−^ dataset) and 4% of NCOs did not match with any DMC1 hotspot (Table 1-table supplement 3). The inferred PRDM9 alleles responsible for the identified NCOs (Figure 4) were fully consistent with the *Prdm9* genotypes of the particular backcross generation of the emerging consomic strains. While the F1 parents were *Prdm9*^B6/PWD^ heterozygous, the *Prdm9*^B6/B6^ homozygosity was achieved no later than at BC3 in all available lines with the obvious exception of the B6.PWD-Chr 17 line (*Prdm9* is localized on Chr 17, Figure 4-table supplement 1). Remarkably, we detected two NCOs initiated by PRDM9^B6^ on Chr 17 (PWD), while only one NCO initiated by PRDM9^PWD^ (Figure 4). Despite the limited sample size, the ratio was in agreement with the previous finding that 71.8% of DMC1 signal occurred in asymmetric DSB hotspots in (PWD × B6)F1 or (B6 × PWD)F1 hybrids^15^. The number of BC generations with the *Prdm9*^B6/PWD^ or *Prdm9*^B6/B6^ genotype corresponded well to the proportion of NCOs initiated by the respective alleles (r = 0.80, P < 0.0001).

**Figure 4.**
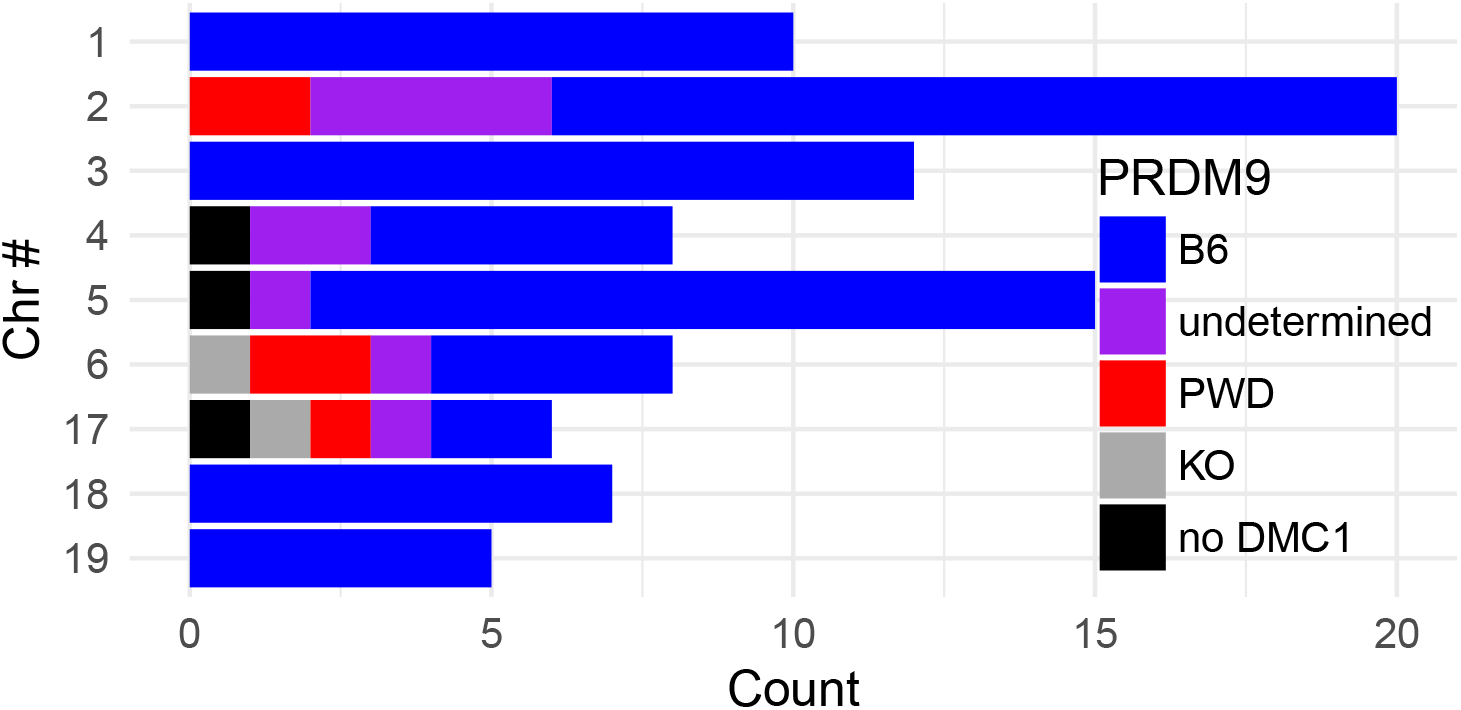
The inferred PRDM9 allele initiating the autosomal DSBs repaired by NCOs. The initiating PRDM9 allele was inferred based on the overlap with parental B6 and PWD strains and the detected PRDM9 binding motif in datasets of Davies and coworkers^15^ and Smagulova and coworkers^34^.

### Chromosome-wide colocalization of NCO and CO events

To compare the localization and properties of NCOs and COs we genotyped 102 (B6 × PWD)F1 male-derived meioses using MegaMUGA genotyping arrays. The identified 1486 autosomal COs were mapped with ~80 kb median resolution sufficient for monitoring the CO spatial distribution at the chromosomal level. Out of the total, 627 (42%) CO sites were found repeatedly (2 to 13 times) within a single interval defined by MegaMUGA, showing the presence of an active CO hotspot.

The COs were highly abundant in the subtelomeric regions and at the 2^nd^ decile of the chromosomal lengths (Figure 5a) in agreement with the non-uniform spatial distribution previously described in male meiosis of mice^36,37^ as well as of humans and other species^38–40^. The spatial chromosome-wide distribution of NCOs was significantly different from the CO distribution (P ≤ 0.0109, see Materials and Methods). In contrast to COs, a significant deficit of NCO events was observed in the 10^th^ decile of the chromosomal length (P ≤ 0.0333). Moreover, NCOs were more abundant around the 4^th^ decile of the chromosomal length (Figure 5b). The NCOs were compared to two unrelated datasets of COs, both showing a difference. First, NCOs were compared to the COs in (B6 × PWD)F1 male meioses detected in this study (P = 0.0029, permutation test) to control for the genetic background. Second, NCOs were compared to a dataset of male and female COs from the G2:F1 population of the Collaborative Cross project^37^ (P = 0.0109, permutation test, see Materials and Methods) to account for the previously described sex differences in the CO distribution. Consistently in both comparisons, the frequency of NCO events in the 10^th^ decile of the relative chromosomal length was lower than that of CO events (P = 0.013, P = 0.0333, respectively, permutation test adjusted for multiple testing). Since none of the published chromosome-wide distributions related to recombination activity including SPO11-oligos in B6^35^, DMC1 heat and H3K4me3 enrichment in B6 and (B6 × PWD)F1^15,34^ had comparable abundance of signal at the subtelomeric region, the resolution of DNA DSBs by COs is more probable than by NCOs at the subtelomeric regions compared to the other parts of the genome. To compare the CO and NCO outcomes and their PRDM9 initiation with a kilobase resolution we focused on a subset of (B6 × PWD)F1 COs whose position determined by the MegaMUGA chip overlapped a single DMC1 (B6 × PWD)F1 hotspot^15,34^ (hereafter DMC1-unique CO sites). We assumed that the respective COs were initiated in the centers of such DMC1 sites. Limiting the DMC1-unique CO sites to those of the inferred PRDM9^B6^ origin, the COs were found more frequently in the NCO sites than in the sites without detected NCOs (P = 0.0401, Mann-Whitney test), suggesting colocalization of CO and NCO hotspots.

**Figure 5.**
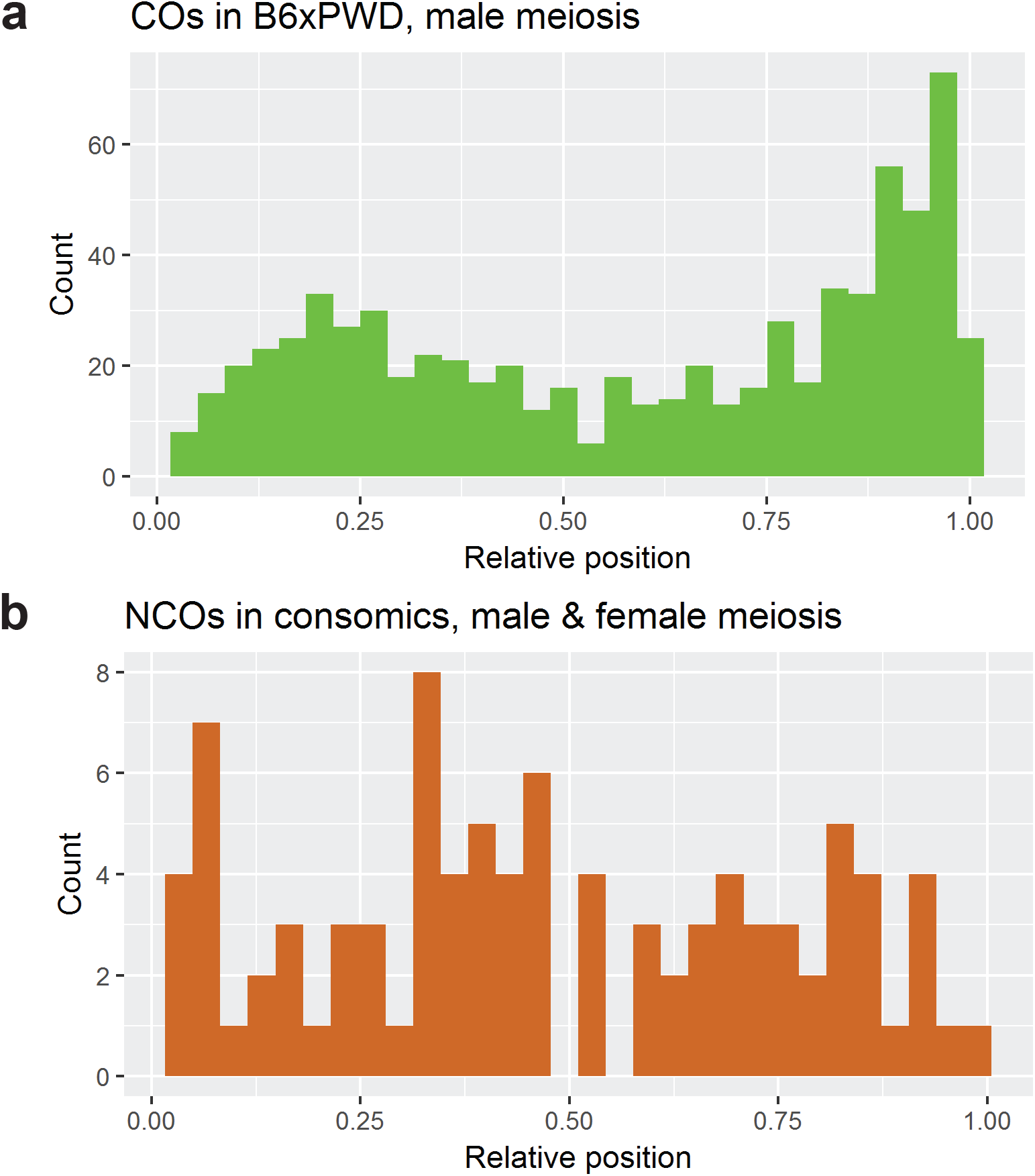
Histograms of relative chromosomal positions of **a** CO and **b** NCO events. The spatial distributions are significantly different, with a significant deficit of NCO events in comparison to CO events at the 10^th^ decile of relative length.

### Higher H3K4me3 and DMC1 hotspot activity in NCO and CO sites

To search for the differences between CO and NCO resolution pathways, we compared histone 3 lysine 4 trimethylation and DMC1 protein abundance at the CO, NCO and DSB sites. The CO sites were filtered to the above-defined subset of DMC1-unique CO sites. NCO sites were filtered to those overlapping DSB sites (DMC1) in the (B6 × PWD)F1 database also used as a resource of DSB sites. We assumed the localization of CO, NCO and DSB sites in the DMC1 site center and we filtered for the sites initiated by PRDM9^B6^ to take into account the differences between the models for CO, DSB and NCO detection. The (B6 × PWD)F1 DMC1 heat was higher in both NCOs (median = 176) and COs (203) than in general DSB (40.4) sites (Figure 5-figure supplement 1a). Similarly, the H3K4me3 enrichment was higher in both NCOs (median = 1.33) and COs (1.70) than in general DSBs (0.46) (Figure 5-figure supplement 1b) but to a lesser extent. The higher DMC1 and H3K4me3 signal in both COs and NCOs is consistent with the higher frequency of the CO and NCO events and thus higher chance of their detection.

The COs, DSBs and NCOs overlapped to a comparable extent with genes (47%, 54%, 54%, respectively). In particular, the overlaps with the gene promoters (3%, 3%, 3%) and exons (3%, 5%, 6%) were comparable as well.

### Asymmetry of PRDM9 binding and deficiency of nonparental NCOs support the DSB repair mechanism of hybrid sterility

During species evolution the PRDM9 hotspots are gradually eroded due to the biased gene conversion that favors the PRDM9 binding sites with lower affinity^3,41–43^. In hybrids between mouse subspecies *M. m. musculus* represented by the PWD mouse strain and *M. m. domesticus* (strain B6) the presence of (partially) erased parental PRDM9 hotspots leads to their heterozygosity. The resulting PRDM9 binding site “asymmetry”^15^ is supposed to cause hybrid sterility of (PWD × B6)F1 males^14,26,27^. It was proposed that the DSBs in asymmetric hotspots are more difficult to repair or are fixed by the noncanonical sister-chromatid repair^15^. The consequent lack of symmetric sites is supposed to result in synaptic defects of homologous chromosomes and male sterility^16,28,44,45^.

If, as proposed, the repair of DSBs by non-sister homologous recombination is inhibited by the asymmetry of their initiation, then a lower than expected frequency of NCOs overlapping the asymmetric DMC1 hotspots could be expected. To examine the idea we classified all NCOs according to the presence of their conversion tract sites within the B6, PWD, (B6 × PWD)F1 or (PWD × B6)F1 DMC1 hotspot datasets^15^ and their *Prdm9* binding motif (Table 1-table supplement 3) Only 17.4% of NCO conversion tract sites (15 out of 86) were classified as “nonparental” (named “novel” in ^34^), being absent in PWD and B6 DMC1 hotspot datasets while present in F1 hybrids (Table 1-table supplement 3, see Materials and Methods for exact definition). This is in contrast with 32% (P = 0.0099, OR = 0.481, 95% CI = (0.266, 0.817), logistic regression, see Materials and Methods) of nonparental DMC1 hotspots observed in (B6 × PWD)F1 and (PWD × B6)F1 hybrids used as a proxy for the asymmetry. An even higher proportion of nonparental DMC1 sites could be expected in the consomic model because the NCOs were found almost exclusively in the sites where PRDM9^B6^ bound to the PWD chromosome. By default, such sites are more likely to be asymmetric. Thus, the observed deficiency of NCOs that have arisen from asymmetric hotspots matched up with the expected difficulties of repair from the homologous chromatid template. In theory, this might be explained by an increased tendency for CO repair in the nonparental sites. However, this alternative is highly unlikely since we found almost identical proportion of nonparental CO (19.0%) and NCO (17.4%) sites. To conclude, the observed deficiency of both, NCOs and COs incurred from asymmetric hotspots in accordance with the expected difficulties of their repair from the homologous chromatid template.

Next, we asked whether the autosomes more sensitive to asynapsis display larger proportions of nonparental NCOs/COs. Interestingly, no such correlation was found (P = 0.5092, Spearman’s ρ = 0.1614, Figure 6) suggesting that the DSBs on short, highly asynaptic chromosomes were not particularly more difficult to repair. Rather, the shorter (more asynaptic) chromosomes contain less DSBs in total, making the selection for the properly repaired DSBs more difficult. This finding supports our model of a threshold number of repaired DSBs necessary for the proper pairing^16,45^.

**Figure 6.**
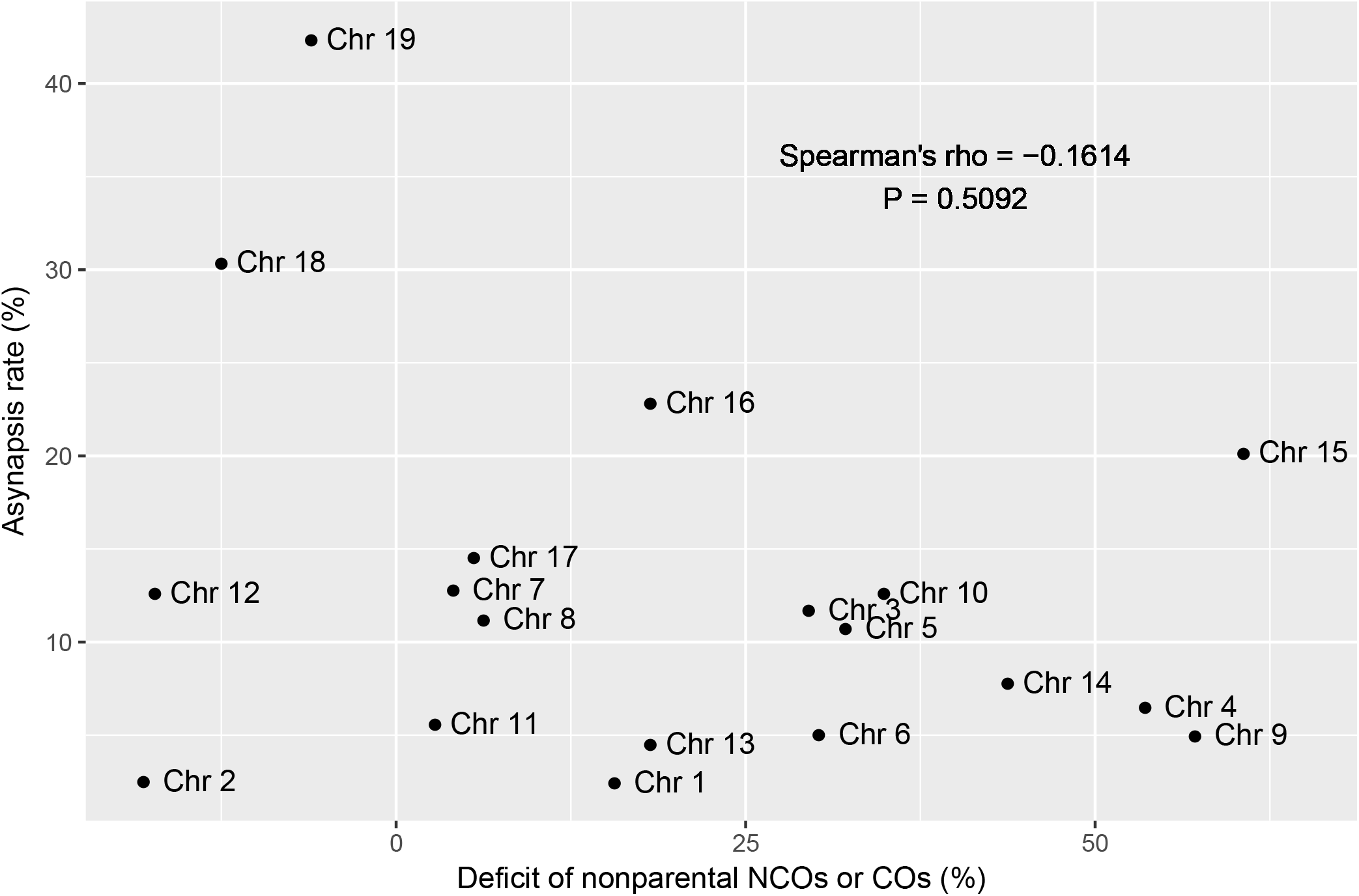
The deficit of nonparental NCOs or COs does not affect F1 asynapsis rates. The NCOs and COs were pooled into one group as their level of nonparental events did not differ. For each autosome the deficit of nonparental NCOs or COs was calculated as the difference between the proportion of nonparental DSBs and the proportion of nonparental NCOs or COs. The asynapsis rate of each autosome was determined for (PWD × B6)F1 males previously^16^. For each autosome the DSBs were paired with NCOs and COs with respect to the (statistically confounding) (B6 × PWD)F1 DMC1 heat determined previously^15^.

### NCO-associated GC bias

GC-biased gene conversion refers to a situation where GC in a conversion tract is transmitted to gametes more often than AT in a conversion tract from GC/AT heterozygotes^46,47^ In our set of NCO sites the bases A or T (so-called “weak” bases, *w*) were converted to bases C or G (“strong” bases, *s*) 3.4 times more often than *vice versa*, showing a GC bias of 77% (P = 2.078 × 10^−8^, two-tailed binomial test, #*w->s* / (#*w->s +* #*s->w*) = 81/(81+24)). The observed GC bias is higher than 68% or 69% reported in humans^30^. Remarkably, we observed a similar GC bias both, in single-SNP NCOs (76%, P = 2.057 × 10^−6^, 42/(42+13)) and multiple-SNP NCOs (“co-conversions”) (78%, P = 9.021 × 10^−5^, 39/(39+11)) (Figure 7a). Our finding is in contrast with the observation in a parallel study^48^ where the *Prdm9*^Hum^-initiated NCOs were analyzed in detail to see the effect of GC bias in *Prdm9*-uneroded genome. The difference may indicate that the mechanisms causing the GC bias operate differently in a *Prdm9*-eroded and *Prdm9*-uneroded genome.

**Figure 7.**
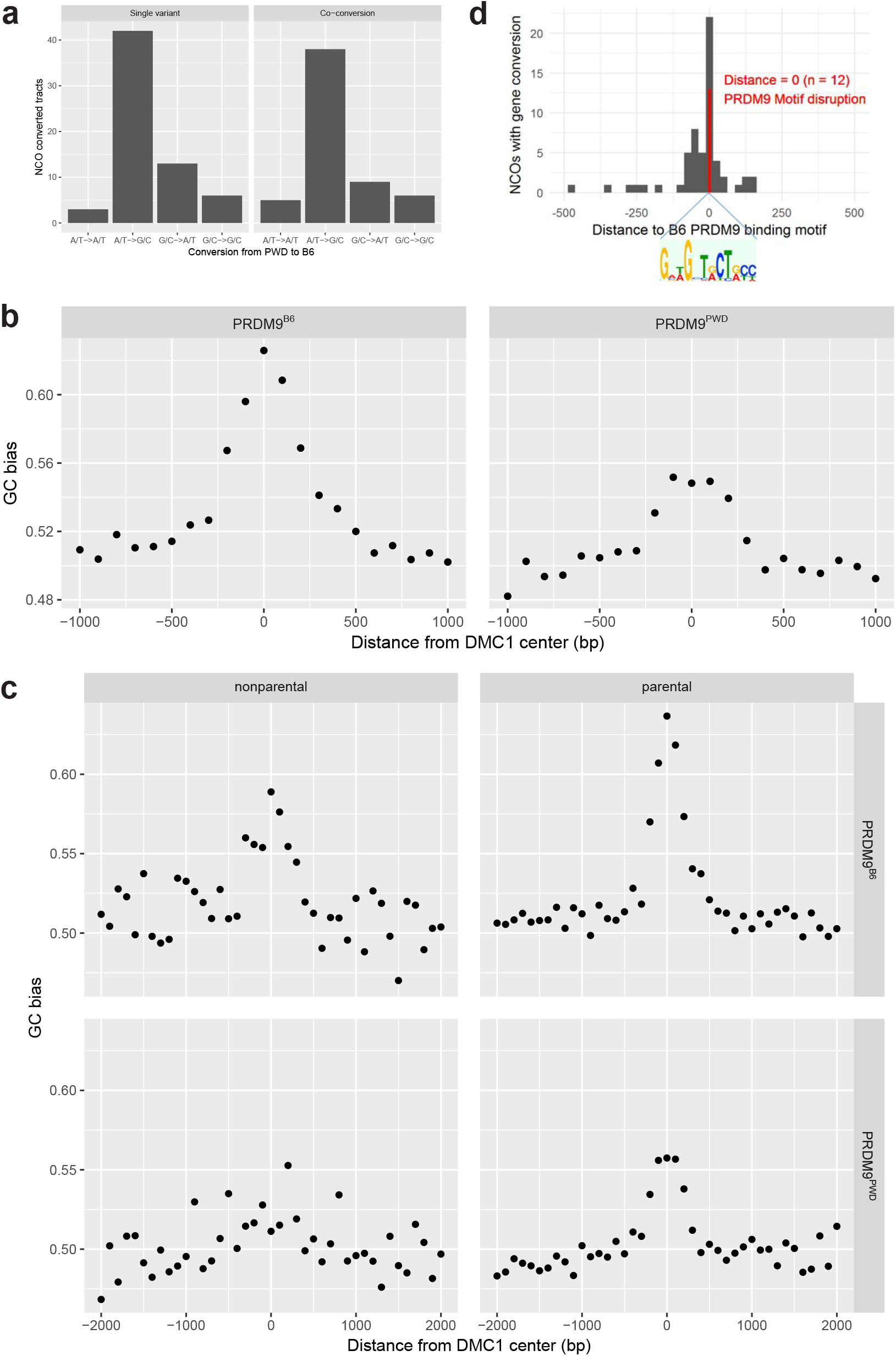
Genome-wide effects of NCO/CO gene conversions. **a** The observed NCO gene conversion converted the A/T “weak” base into the G/C “strong” base 3.4 times more frequently than *vice versa*, the “strong” base into the “weak” base. **b** GC bias already present due to historical recombination in (B6 × PWD)F1 hybrids in DMC1 sites^15^ and ± 1000 bp. The genome is GC-biased (and *Prdm9*-eroded) in the autosomes of both PWD and B6 strains. Each dot represents the average GC bias in the sliding window of 100 bp. **c** The CG bias (shown here for PRDM9^B6^ and PRDM9^PWD^) is higher for parental than for nonparental DMC1 sites. The GC bias was calculated analogously as in (b). **d** Histogram of distances of minimal possible NCO tracts from the inferred PRDM9^B6^ binding motif. NCO gene conversions disrupted the binding motif in 20.6% of cases.

We observed the same pattern of GC bias in the surrounding non-converted sites, both upstream and downstream from the NCO converted sites, most probably as a consequence of historical meiotic gene conversions. As expected, the rate of GC bias decreased with the distance from the *Prdm9* binding sites with no GC bias in the 500 bp distance (Figure 7b). This might be explained by combination of more localized NCO gene conversions and more spread CO gene conversion events occurring in the same site in the history of the locus. In addition, we observed a stronger GC bias for parental (63% for *Prdm9^B6^*, Figure 7c) than for nonparental sites (59%) suggesting that the process of gene conversion leading to GC bias lasted longer in evolution in parental sites and has affected more sites in the surroundings of the PRDM9 motif. Measured alternatively by the level of asymmetry, we also observed a stronger GC bias for the symmetric sites than for the asymmetric ones as expected. Simultaneously, we found that there was an excess of PWD-B6 SNPs in the centers (± 18 bp) of nonparental DMC1 sites (47.6%) than in the centers of parental DMC1 sites (30.0%). These findings are consistent with the view that the binding motifs of nonparental DMC1 sites were disrupted at early generations in the evolution, the binding sites were not used so frequently or even at all since that moment and the gene conversions accompanying the repair of meiotic DSB at these sites created a smaller GC bias. Consistently, the parental DMC1 sites are still active in the parental strains and have created a higher GC bias. Finally, the identified NCO gene conversions disrupted the PRDM9 motif in 20.6% of cases (Figure 7d) showing perfect agreement with the motif erosion localized in a single *A3* hotspot^17^.

## Discussion

In mice, NCOs are the prevalent outcome of the repair of programmed meiotic DSBs with the NCOs/COs overall ratio 9:1, based on cytological data and sperm typing^7,49,50^. Despite their prevalence and assumed functional significance for meiotic synapsis of homologous chromosomes^10,11,50^ little is known about their genome-wide distribution and properties. In this study, we used whole-genome sequencing to detect and characterize a unique dataset of 95 NCO events in 10 mouse chromosomes using the B6.PWD-Chr # panel of mouse consomic strans. To compare the chromosome-wide distribution and colocalization of NCOs and COs we identified nearly 1500 CO events resulting from (B6 × PWD)F1 meioses. When writing the manuscript another genome-wide study of mouse NCOs was reported^48^. Previous studies of individual hotspots in mice and humans determined the average length of CO-associated conversion tracts to be ~500 bp^17,18,50^. The NCO gene conversion tracts were much shorter, 50–300 bp at the mouse *Psmb9* hotspot and two hotspots on chromosome 1 ^51,52^ and 55–290 bp at the MHC hotspot in human chromosome 6 ^18^. More recently, a human genome-wide study based on Illumina SNP arrays indicated the multiple-SNP NCO gene conversion tract length in the range of 100–1000 bp^53^, while most of NCO events involved only one SNP due to low SNP density. Thus our estimate, 32 bp, based on the screen of 10 mouse chromosomes is much shorter than reported previously for individual hotspots but close to another recent mouse study showing 30 bp and 41 bp for two different *Prdm9* alleles^48^. Another difference from the human studies is an almost complete absence of complex NCO gene conversions in our dataset. Only one out of 94 NCO events was a 231 bp long complex conversion interrupted by one nonconverted SNP in contrast to 46% or 65% of complex events found in human studies^30,53^.

Until recently, the studies of NCOs in mice were based on the analysis of individual recombination hotspots^7,17,51,52,54^. Our finding of 96% colocalization of NCO conversion tracts with DMC1 hotspots confirmed both the origin of NCOs from meiotic DSBs and our efficiency to filter out the false positive events. Moreover, the almost exclusive occurrence of NCO events within DMC1 hotspots testifies against a significant share of female-specific NCOs in our dataset. The DMC1 data used for comparison came exclusively from the males, while the identified NCOs were of male and female origin. The first study measuring the DMC1 signal in female meiosis^55^ showed only negligible sex difference in the genomic sites of DMC1 signal. What differed was the activity of DMC1 hotspots corresponding to the variations in H3K4 trimethylation and affected in males by DNA methylation. Thus the findings by Brick and coworkers^55^ of negligible sex differences in the localization of DSBs can be extended to NCOs.

Based on cytological and sperm typing studies the NCOs are supposed to represent ~90% of all repaired DSBs; however the ratio varies depending on the particular hotspot^49,52^. The mechanism responsible for the decision between CO and NCO resolution is largely unknown. Recently, a study based on whole-genome sequencing of single sperm indicated that PRDM9 protein binding to the uncut template homolog decreases the time necessary for homolog recognition and single-strand DNA invasion into the duplex DNA and increases the chance to process the recombination intermediate into CO^56^. The same study confirmed the previous findings of nonrandom distribution of COs along chromosomes in male meiosis of mice and humans, showing CO excess in subtelomeric regions and their paucity at the centromeric ends^36–38,49,52^. Our findings of the excess of COs in the subtelomeric regions and in the second decile of the chromosomal length in (B6 x PWD) male meiosis and significant deficit of NCOs in the 10^th^ decile of relative chromosome length are supported by these studies and show that DNA DSBs at the subtelomeric regions are more prone to be resolved as COs relative to NCOs.

NCOs are associated with biased gene conversion (BGC), which results in uneven recovery of each allele in the next generation and can be generated by recombination initiation bias when the PRDM9 binding motif of the recipient haplotype has higher probability to be targeted by SPO11 than the donor (template) haplotype. The mechanism of GC-biased gene conversion (gBGC) can be explained by biased repair of mismatched AT/GC alleles in heteroduplex DNA favoring strong G/C over weak A/T base pairs^46,48^ or subtle preference of the A/T allele at initiation of recombination^48,57^. Meiotic drive in favor of the GC allele (~68%) was documented in the human genome^30,53,58,59^ and more recently in mice (60%-68%)^48^. In our study a significant GC bias (76%) was detected genome-wide in the NCO sites, but also in the surrounding non-converted sites, suggesting the DSB repair activity in the past. The GC bias was gradually decreasing outwards from the DMC1 center, with no bias in the ±500 bp distance, similarly as reported earlier^47^. This fading effect most likely resulted from the historical recombination accompanied by NCO and CO gene conversions, which had migrated from the DSB initiation site. The humanized *Prdm9^Hum^* binding sites have no history of erosion in classical (e.g., C57BL/6J) or wild-derived laboratory (e.g., PWD/Ph, CAST/EiJ) mouse genomes and the surrounding regions are not GC-biased. Li and coworkers^48^ observed the differences between NCO gene conversion tracts initiated by PRDM9^Hum^ (41 bp) and PRDM9^Cast^ (mean = 30 bp) in the genomes of B6 and CAST crosses. Here, we showed the mean conversion tract length of 32 bp, identical as for PRDM9^Cast^, for the NCO sites almost exclusively initiated by PRDM9^B6^. We suggest a possible relationship between the degree of PRDM9 binding site erosion and the length of a gene conversion tract that is not based on inherent properties of the *Prdm9* alleles. Rather, the mean length of conversion tracts could reflect the preceding activity of PRDM9 in the binding site and surrounding regions and consequent GC bias. Such GC-biased regions are significantly more resistant to be converted again, leading to shorter gene conversion lengths. Consequently, this could imply a gradual decrease in time of building the GC bias as the surrounding bases of the PRDM9 binding site become increasingly G/C rich, which limits the tract length. Asymmetric binding of PRDM9 to allelic sites was identified in several combinations of intersubspecific F1 hybrids^15,34,42^. It was proposed that excessive asymmetry of DNA DSB hotspots is the key feature of early meiotic arrest in the (PWD × B6)F1 model of hybrid sterility^15^. Although many aspects of the role of *Prdm9* in hybrid sterility remain to be clarified^60^, new data bring additional support for the DSB asymmetry hypothesis^16,45,56^. The repair of asymmetric DSBs can be delayed or aborted, resulting in chromosome breakage and their deficiency in hybrids. Indeed, only 12% of our NCOs appeared as nonparental, generated from a subset of asymmetric DSBs. Despite being significantly below the expected 32%, it shows that some asymmetric sites are either only partially eroded and can be distinguished in a fraction of cells, or the sister chromatid taboo is breached and sister chromatids are used as a template, or a combination of both.

## Materials and Methods

### Animals and ethics statement

The animals were maintained at the Institute of Molecular Genetics in Prague, Czech Republic. The project was approved by the Animal Care and Use Committee of the Institute of Molecular Genetics ASCR, protocol No 005/2017. The principles of laboratory animal care, Czech Act No 246/1992 Sb., compatible with EU Council Directive 86/609/EEC and Appendix of the Council of Europe Convention ETS, were observed. The PWD/Ph inbred strain originated from a single pair of wild mice of the *M. m. musculus* subspecies trapped in 1972 in Central Bohemia, Czech Republic^21^. The C57BL/6J (B6) inbred strain was purchased from The Jackson Laboratory. The panel of 27 chromosome substitution strains C57BL/6J-Chr #PWD (B6.PWD-Chr #) maintained by the Institute of Molecular Genetics AS CR, Czech Republic, was prepared in our laboratory^20^. All mice were maintained in a 12 hr light/12 hr dark cycle in a specific pathogen-free barrier facility. The mice had unlimited access to a standard rodent diet (ST-1, 3.4% fat, VELAZ) and water. All males were sacrificed at age 40-60 d. The tails of progenitors of chromosomal substitution strains have been archived in a - 80 °C freezer.

### NGS sequencing and genotyping

Partially purified DNA from mouse spleens was isolated by Puregene Core Kit A (QIAGEN 158267) according to the manufacturer’s instructions. Next generation sequencing was performed in Illumina HiSeq X to obtain paired-end reads of lengths 2×150bp. The Kapa library preparation protocol was used. The obtained reads were checked for quality by FASTQC and deduplicated. The paired-end reads were aligned to both mm10 reference genome and to PWK pseudo-reference genome and used for identification of NCO sites as described below.

### Detection of noncrossover sites

The NCO sites were identified by subsequent filtering steps using both the conservative and anti-conservative method. In the conservative method, the Illumina paired end 2×150 bp reads were aligned to both B6 reference and PWD pseudo-reference fasta files. The candidate reads were required to align perfectly to the B6 reference and not perfectly but at most with six variants to the PWD pseudo-reference. The detected variants in the reads were then compared to four datasets of Illumina reads of B6, PWK, PWD^15,61^ and the B6.PWD-Chr # strains (this study) and the sites heterozygous either in B6 or PWD parental strains were excluded. Similarly, the gene conversion candidate sites in the transferred PWD chromosomes of B6.PWD-Chr # strains were required to be homozygous.

Further, two regions of B6 origin of lengths ~3Mb were found on the background of transferred, otherwise PWD chromosomes in the B6.PWD-Chr 3 and B6.PWD-Chr 4 strains, respectively: chr3:22181981-25498721 and chr4:153634374-156356368. Most likely, they resulted from hidden double crossovers during the preparation of chromosomal substitution strains. As such they were excluded from further analyses.

Using the NGS data, all sites were manually curated using the IGV genome browser. Other steps of validation were performed subsequently and independently, namely Sanger sequencing, overlap with the known sites of recombination detected in previous studies^15,34^ as described below.

### Noncrossover site amplification and Sanger sequencing

The DNA from mouse tails was amplified at selected NCO sites (see primer list). Samples were amplified by a BIOER XP Cycler with program: 95 °C for 2 min followed by 40 cycles at 95 °C for 10 s, 55 °C for 20 s and 72 °C for 30 s, and single step at 72 °C for 3 min. Samples showing a single correct size band were treated by ExoSAP-IT enzyme according to manufacturer’s instructions (Thermofisher Scientific 78201) and sequenced.

### *Prdm9* exon 12 amplification

DNA was amplified with primers Exon12-L1 – TGAGATCTGAGGAAAGTAAGAG and Exon12-R – TGCTGTTGGCTTTCTCATTC with PCR program: 94 °C for 5 min and 40 cycles at 94 °C for 30 s, 61 °C for 1 min, 68 °C for 2 min, and a single step at 72 °C for 7 min. The PCR products were resolved in 2% agarose gel allowing visual discrimination of B6 from PWD alleles of *Prdm9* by their different migration.

### Dynamics of NCO gene conversion occurrence

For inferring the dynamics of gene conversion during 10 generations of backcrosses, the converted SNPs in NCO sites were additionally sequenced by Sanger sequencing in a step-wise procedure. First, the male parents from BC-10 and BC-6 generations, then the samples from the remaining generations between BC-1, BC-6 and BC-10 were either genotyped, when the closest known generation before was PWD/B6 and the closest generation after was B6/B6, or inferred when the closest known generation before and after were both of the same genotype. The parents from BC-2 and BC-3 used for preparation of B6.PWD-Chr 2 were not archived.

### Overlap of NCOs and COs with DMC1 and H3K4me3 ChIP-seq sites

To determine the overlap of NCO and CO sites with DMC1 hotspots, we used the DMC1 datasets of (B6 × PWD)F1, (PWD × B6)F1, B6 and PWD males from the previous studies^15,34^. The datasets differed in the number of detected hotspots between the biological replicates^15, 34^; however, the extra DMC1 hotspots had the lowest heat. The heat of DMC1 sites overlapping between the biological replicates was highly correlated (ρ = 0.92, P < 10^−16^, for (B6 × PWD)F1). Importantly, the extent of the overlap of autosomal NCO sites with the respective DMC1 datasets differed by only 2% (higher in Davies and coworkers ^15^). Furthermore, only the latter datasets ^15^ provide the information about the inferred PRDM9 initiation of each DSB and the level of asymmetry. For the Chr X, which is missing in the dataset of Davies and coworkers ^15^, we used the information on the DMC1 overlap from datasets of Smagulova and coworkers ^34^. Similarly, the vast majority (98%) of NCOs overlapping either (B6 × PWD)F1 or (PWD × B6)F1 DMC1 sites overlapped each other. The DMC1 sites were considered uniformly of 2001 bp length, consistently with the reported median DMC1 site length^34^. Because the autosomal COs in our study were detected in the (B6 × PWD)F1 hybrids, we assumed their localization in DMC1 hotspots of the (B6 × PWD)F1 DMC1 dataset. To investigate the presence of H3K4me3 signal in NCO and CO sites of (B6 × PWD)F1, (PWD × B6)F1 and B6 males we used the data from Davies and coworkers ^15^. For the comparison of the amount of DMC1 and H3K4me3 signal in NCO and CO sites, we used only the data from the (B6 × PWD)F1 dataset^15^. The H3K4me3 sites were considered uniformly of the 2001 bp length.

### Comparison of NCO and CO spatial chromosome-wide distribution

The NCOs were compared to the COs in (B6 × PWD)F1 male meioses generated in this study to control for the genetic background. Next, the NCOs were compared to both male and female COs inferred from the G2:F1 population of the Collaborative Cross (CC) project^62^. In both cases, the comparison was performed by a permutation test. The respective CO events were binned into 10 groups according to their relative chromosomal coordinate to give the estimates of the relative proportions of CO events in each bin, which were used to specify the probabilistic distribution. Provided the null hypothesis NCOs do not differ from COs, a dataset of 91 NCO events was simulated from the respective CO distributions. The simulations were performed 10,000 times and the real NCO distribution was compared to the expected NCO distribution. In addition, to compare with the male and female COs from the CC project, the CO distribution based on 50% of male COs and 50% of female COs and CO distribution based on 25% of male COs and 75% of females COs were calculated as two extreme possibilities. This was done to account for the uncertainty of sex for which the NCOs were observed. The more conservative p-value was reported.

### Proportion of nonparental NCO, CO and DSB events as a proxy for the level of asymmetry

We compared the abundance of nonparental NCO, CO and general DSB sites to assess the hypothesis that the asymmetric DSB sites are harder to repair and are possibly repaired by sister chromatids. As a result, they would not be observed as either NCO or CO outcome. Here we defined an “event“ as either an NCO event, CO event, or meiotic DSB event. Second, we defined the NCO/CO/DSB events as “nonparental“ if found in the DMC1 dataset of (B6 × PWD)F1 and/or (PWD × B6)F1 hybrids and not in the DMC1 dataset of B6 or PWD parents and B6.*Prdm9*^−/−^. Third, the “parental“ events were defined as found in the DMC1 dataset of (B6 × PWD)F1 and/or (PWD × B6)F1 hybrids and also in one of the B6 or PWD DMC1 parental datasets but not in the B6.*Prdm9*^−/−^ dataset. The events not found in any of (B6 × PWD)F1 or (PWD × B6)F1 DMC1 datasets and the events found in the B6.*Prdm9*^−/−^ DMC1 dataset were excluded from the analysis. We calculated the proportion of nonparental events as a ratio (# of nonparental events) / (# of nonparental and parental events) for the NCO events. For the comparison to CO and DSB events we calculated the ratio similarly, but considered two additional features of the DMC1 data. First, DMC1 heat was significantly correlated with the level of asymmetry (ρ = −0.26, P < 0.0001; ρ = −0.22, P < 0.0001), and with the proportion of nonparental sites (ρ = 0.11, P < 0.0001; ρ = 0.08, P < 0.0001), in (B6 × PWD)F1 hybrids for both PRDM9^B6^ and PRDM9^PWD^ initiated sites respectively. In addition, the median of DMC1 heat was much lower in the overall DSB hotspots than in the DSB hotspots with the detected NCO/CO occurrence (Figure 5-figure supplement 1a). Thus, the DMC1 heat statistically confounded the relationship between the parental/nonparental origin of the events and the class of the event (NCO/CO/DSB) and was controlled for (Table 1-figure supplement 1) in the used logistic regression models. Second, a vast majority (~90%) of the detected NCO events were initiated by PRDM9^B6^ and all the detected NCO events occurred on the PWD chromatid. This was an inherent consequence of using chromosomal substitution strains as a model. This means that the NCO events occurred by design in more asymmetric sites than the DSB sites detected in (B6 × PWD)F1 and (PWD × B6)F1 hybrids by DMC1 ChIP-seq. Thus, the referred difference of proportions of nonparental NCO and nonparental DSB events should be interpreted as a conservative (minimal) estimate implying that the deficit of nonparental NCO events is likely to be even larger.

### Statistics

All statistical tests were performed and figures were plotted using R 3.4.4, the tidyverse package (dplyr, ggplot)^63^ in particular. The mean lengths of NCO gene conversions were assumed to follow an exponential distribution according to Li and coworkers^48^ to provide a direct comparability. Composite likelihood of the rate parameter λ was estimated using the probabilities that the neighboring SNPs were converted. Other, non-standard statistical analyses are described in the Materials and Methods part above. Uniformly, the permutations tests used to calculate the expected overlap between the genomic features were done using N = 10,000 permutations. The standard statistical tests used are mentioned in the Results part.

## Data availability

The NGS data are deposited under SRA BioProject number PRJNA564925.

---

**Figure 5–figure supplement 1.**
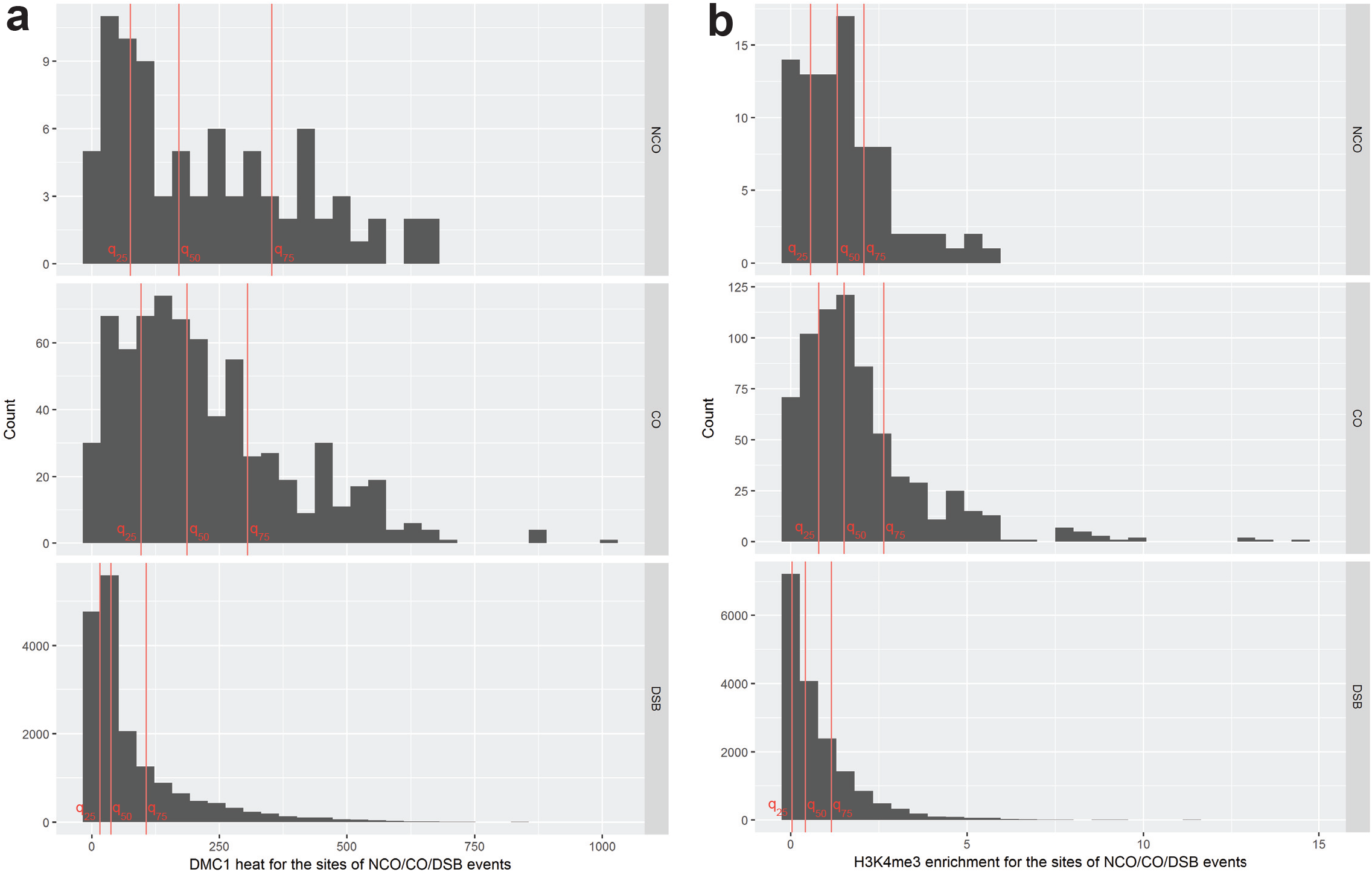
Histograms of the DMC1 heat and H3K4me3 enrichment in (B6 × PWD)F1 hybrids. **a** DMC1 data from ^64^. Red vertical lines labeled q_25_, q_50_, q_75_ represent 25^th^, 50^th^ and 75^th^ quantiles of the respective distributions. **b** H3K4me3 data from ^64^. Red vertical lines labeled q_25_, q_50_, q_75_ represent 25^th^, 50^th^ and 75^th^ quantiles of the respective distributions. 1 CO (0.14%) and 68 DSBs (0.39%) with H3K4me3 enrichment greater than 15 are not visualized.

**Table 1–figure supplement 1.**
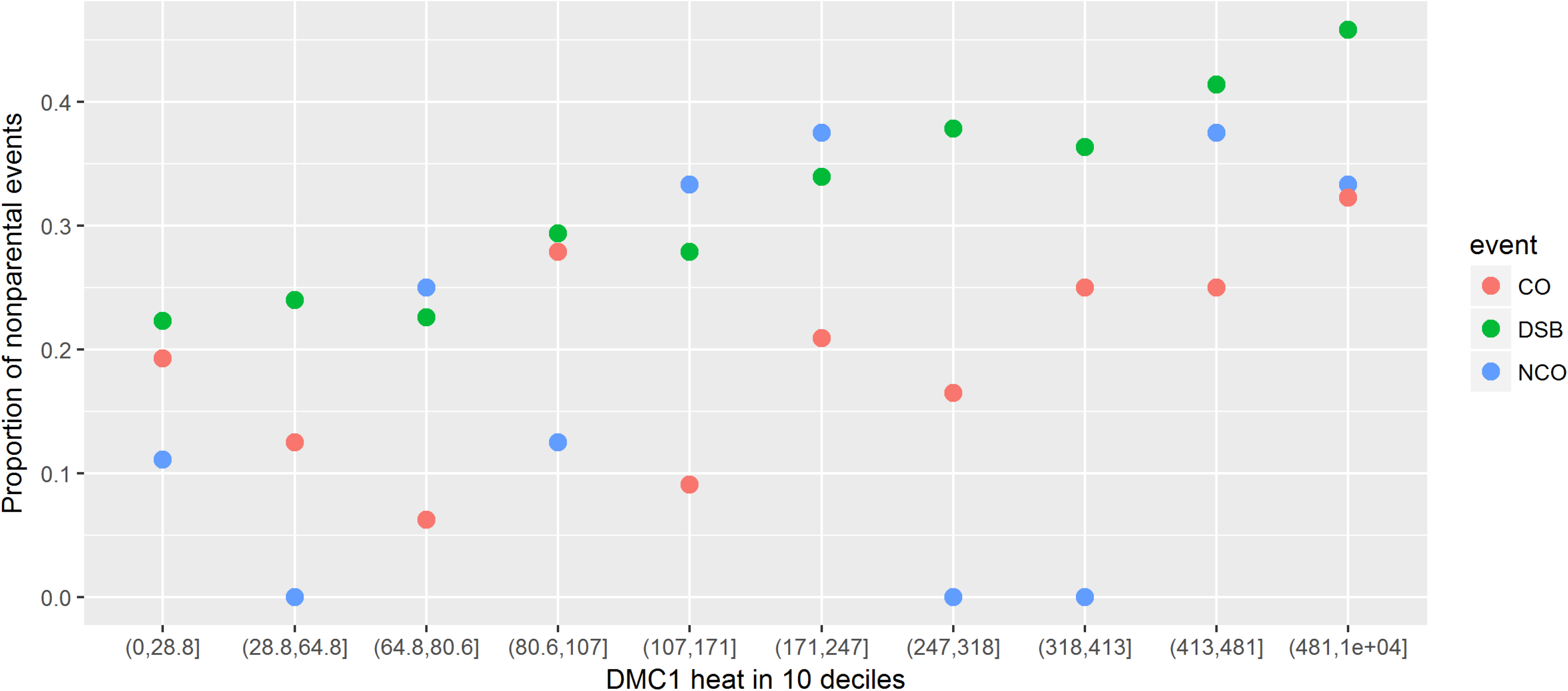
Deficiency of nonparental NCO and CO sites. The nonparental NCO and CO sites are deficient in comparison to total nonparental DSB sites. The DMC1 heat is binned into deciles according to DMC1 heat for NCO dataset. This was done to visually account for the statistically confounding DMC1 heat. The DMC1 heat data is from ^64^.

**Table 1–table supplement 1.**
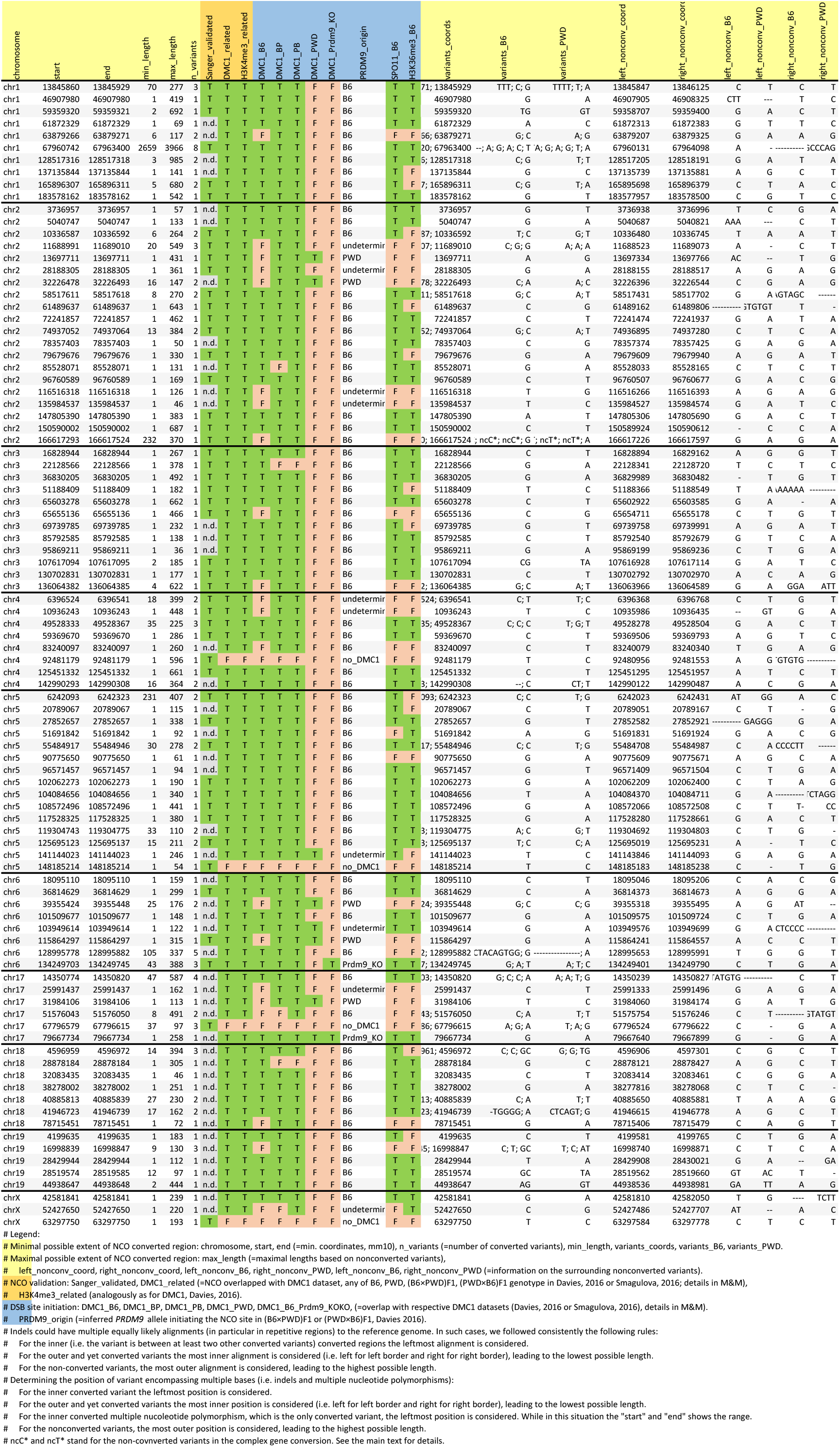

**Table 1–table supplement 2.**

| Chromosome | Length_mm10 | SPO11_oligos | NCO_count |
| --- | --- | --- | --- |
| chr1 | 191909192 | 69606 | 10 |
| chr2 | 178326651 | 73115 | 20 |
| chr3 | 156398855 | 53878 | 12 |
| chr4 | 152055611 | 60332 | 8 |
| chr5 | 147919674 | 62217 | 15 |
| chr6 | 146336543 | 52717 | 8 |
| chr17 | 91707462 | 39641 | 6 |
| chr18 | 87452634 | 34544 | 7 |
| chr19 | 58205856 | 25629 | 5 |

**Table 1–table supplement 3.** Classification of NCOs based on the inferred initiating PRDM9 allele and predicted hotspot symmetry.

| Predicted NCO class | Parentality | DMC1 dataset * |  |  |  | Hotspot origin * | No of NCOs |
| --- | --- | --- | --- | --- | --- | --- | --- |
|  |  | PWD | B6 | B6×PWD | PWD×B6 |  |  |
| Weak PRDM9 motif, symmetric | parental | + | + | + | + | undetermined | 2 |
| PWD sym | parental | + | - | + | + | PWD | 5 |
| PWD asym | nonparental | - | - | + | + | PWD | 0 |
| B6 sym ** | parental | - | + | + | + | B6 | 66 |
| B6 sym | parental, incomparable | - | + | - | - | B6 | 2 |
| B6 asym † | nonparental | - | - | + | + | B6 | 10 |
| Weak PRDM9 motif, asymmetric † | nonparental | - | - | + | + | undetermined | 8 |
| Anomalous, default ‡ | anomalous | + | + | + | + | B6.*Prdm9*<sup>-/-</sup> | 2 |
| Anomalous, DMC1 negative | anomalous | - | - | - | - | none | 4 |

\* Davies et al. 2016, Smagulova et al. 2016
\*\* includes 1 NCO from (PWD-, B6+, B6×PWD- and PWD×B6+)
† includes 1 NCO from (PWD-, B6-, B6×PWD- and PWD×B6+)
†† includes 1 NCO from (PWD-, B6-, B6×PWD- and PWD×B6+)
‡ includes 1 NCO from (PWD-, B6+, B6×PWD+, PWD×B6+)

**Figure 4–table supplement 1.**
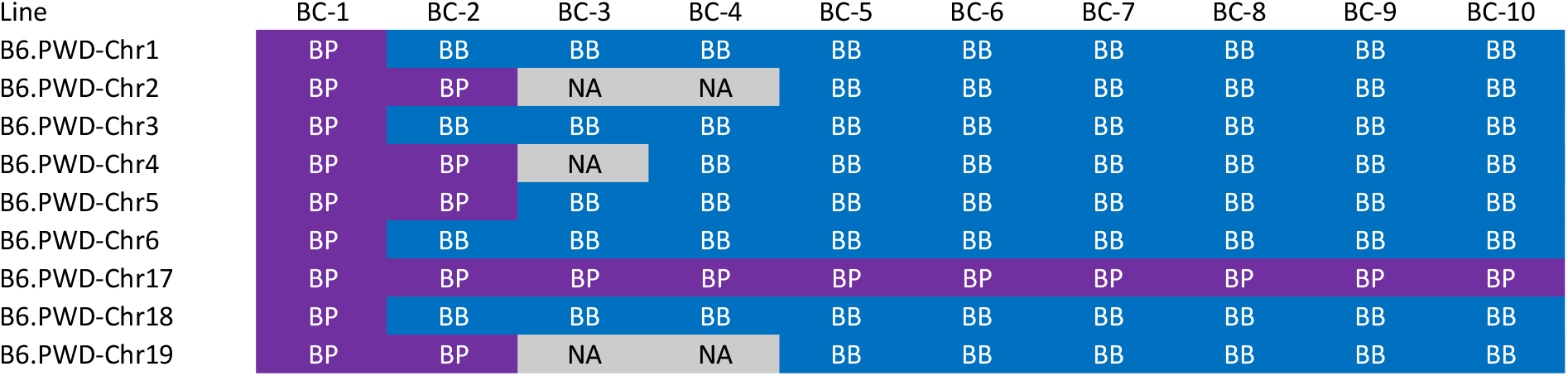
Genotype of *Prdm9* in 10 generations of backcrosses used for preparation of chrosomal substitution strains.

